# The critical role of the ventral temporal lobe in word retrieval

**DOI:** 10.1101/2021.11.18.469030

**Authors:** Kathryn M. Snyder, Kiefer J. Forseth, Cristian Donos, Patrick S. Rollo, Simon Fischer-Baum, Joshua Breier, Nitin Tandon

**Affiliations:** Vivian L. Smith Department of Neurosurgery, McGovern Medical School at UT Health Houston, Houston, TX, USA; Texas Institute for Restorative Neurotechnologies, The University of Texas Health Science Center at Houston, Houston, TX, USA; Faculty of Physics, University of Bucharest, Bucharest, Romania; Department of Psychological Sciences, Rice University, Houston, TX, USA; Memorial Hermann Hospital, Texas Medical Center, Houston, TX, USA

**Keywords:** dysnomia, intracranial EEG, language, lesion-symptom mapping, temporal lobe epilepsy

## Abstract

Deficits in word retrieval are a hallmark of a variety of neurological illnesses spanning from dementia to traumatic injuries. The role of the dominant temporal lobe in fluent naming has been characterized by lesional analyses, functional imaging, and intracranial recordings, but limitations of each of these measures preclude a clear assessment of which specific constituent of the temporal lobe is critical for naming. We studied a large cohort of patients undergoing surgical resections or laser ablations of the dominant temporal lobe for medically intractable epilepsy (n=95). These techniques are exceedingly effective for seizure control but often result in language declines, particularly in confrontation naming, which can be socio-economically disabling. We used a multivariate voxel-based lesion symptom mapping analysis to localize brain regions significantly associated with visual object naming deficits. We observed that posterior inferior temporal regions, centered around the middle fusiform gyrus, were significantly associated with a decline in confrontation naming. Furthermore, we found that the posterior margin of anterior temporal lobectomies was linearly correlated to a decline in visual naming with a clinically significant decline occurring once the resection extended 6 cm from the anterior tip of the temporal lobe. We integrated these findings with electrocorticography during naming in a subset of this population and found that the majority of cortical regions whose resection was associated with a significant decline overlapped with regions that were functionally most active prior to articulation. Importantly, these loci coincide with the sites of susceptibility artifacts during echo-planar imaging, which explains why this region has not previously been implicated. Taken together, these data highlight the crucial contribution of the posterior ventral temporal cortex in lexical access and its important role in the pathophysiology of anomia following temporal lobe resections. Surgical strategies, including the use of laser ablation to target the medial temporal lobe as well as microsurgical approaches, should attempt to preserve this region to mitigate postoperative language deficits.

## Introduction

Lexical retrieval is the process of extracting a specific phonological form from a stored lexical concept. The lexicon is a theoretical construct, conceived as a hub connecting semantic systems to language-related systems such as phonology and orthography.^1,2^ Disruptions in lexical retrieval result in the “tip-of-the-tongue” (TOT) phenomenon, or failure to retrieve a familiar word with partial recall of features associated with the target word.^3^ While the TOT state occurs occasionally in healthy individuals, it is a hallmark of pervasive anomia following a variety of different brain injuries.^4^

Anomia is particularly prevalent in patients with left language-dominant temporal lobe epilepsy, and temporal lobe resections for seizure control further increase the risk of these naming deficits.^5,6,7,8,9^ However, the precise substrates responsible for this cognitive loss are unclear. In clinical literature, a prominent focus has been on preserving the superior temporal lobe to prevent dysnomia.^10^ As such, a clear understanding of the most critical constituents of lexical retrieval might influence the design of surgical strategies to minimize language declines.

A variety of different methods have previously been used to isolate brain regions essential to lexical retrieval, including lesion deficit mapping,^11,12^ functional imaging,^13,14,15,16,17,18,19^ and non-invasive EEG.^20^ Voxel-based lesion symptom mapping (VLSM) is used to localize brain function on a voxel-by-voxel level and demonstrates the relationship between damage at any given voxel and performance on a behavioral task related to a particular cognitive function.^21^ Traditionally, VLSM analyses use a mass-univariate approach, where the lesion-deficit relationship is determined one voxel at a time. However, this approach fails to incorporate information regarding the spatial relationship between voxels, which assumes that voxels are statistically independent and subsequently leads to a loss of statistical power and a potential bias in localization of significant regions. Using a multivariate VLSM approach mitigates these errors by modeling contributions of multiple voxels simultaneously.^22^ Furthermore, VLSM analyses are confounded by the fact that data are typically collected only after the occurrence of the lesion. Thus, a dataset in which both preoperative and postoperative performance measurements as well as imaging are obtained and in whom controlled lesions are produced are of particular value in evaluating the role of a given region.

We applied multivariate VLSM analysis to a large cohort of patients who underwent surgery in the dominant left temporal lobe for treatment of drug-resistant epilepsy to specifically evaluate which component of the temporal lobe is most critical for naming. We quantified the association between each component of the temporal lobe and the change in picture naming performance to isolate brain regions associated with a decline in visual naming ability. Additionally, we integrated these results with electrocorticography (ECoG) recordings during picture naming performed in a subset of patients who underwent intracranial electrode implantation to localize epileptic foci. This allowed us to co-localize and correlate lesion-deficit findings with brain activity.

## Material and Methods

### Study Population

One hundred and eighty-nine patients (105 females, 5-73 years) underwent surgical resection or ablation in the left temporal lobe for drug-resistant focal epilepsy. Study design was approved by the University of Texas Health Science Center’s committee for the protection of human subjects. Of these, subjects younger than 16 years were excluded (n=7). Left-hemispheric language dominance was confirmed in 146 patients by intra-carotid sodium amytal injection,^23^ fMRI laterality index,^24,25^ or direct cortical stimulation.^10^ Patients with right (n=15), bilateral (n=3), or inconclusive (n=25) language dominance were excluded. Additionally, subjects who were not fluent English speakers (n=3), who had an IQ below 67 (n=4), or who had large structural abnormalities or previous temporal lobe surgeries (n=11) were excluded. Of the remaining 122 patients, 95 patients (53 females, 17-73 years, mean FSIQ 97 +/- 15) underwent neuropsychological testing and MRI prior to and following surgery. The other 27 subjects did not have all data points and were excluded. Neuropsychological testing sessions included the full Boston Naming Test (BNT), which consists of 60 black and white line drawings of objects that subjects must name within 20 s. ^26^ Seizure outcomes were reported one year following surgery using the International League Against Epilepsy (ILAE) scoring system with scores of 1 (n=57), 2 (n=6), 3 (n=17), 4 (n=13), or 5 (n=2).

### MRI Acquisition

MRI scans were obtained prior to and following surgery. Preoperative scans were obtained using a 3 T whole-body magnetic resonance scanner (Philips Medical Systems) fitted with a 16-channel SENSE head coil. Images were collected using a magnetization-prepared 180° radio-frequency pulse and rapid gradient-echo sequence with 1 mm sagittal slices and an in-plane resolution of 1 mm isotropic. Postoperative scans were acquired with the same parameters as preoperative scans. Pial surface reconstructions were computed with FreeSurfer (v6.0)^27^ and imported into AFNI (https://afni.nimh.nih.gov). ^28^

Functional MRI was acquired in 35 of these patients (19 females, 19-73 years). Functional images were obtained using a gradient-recalled echo-planar imaging sequence with 33 axial slices of 3 mm thickness and in-plane resolution of 2.75 mm x 2.75 mm (TE = 30 ms, TR = 2,015 ms, flip angle = 90°). Stimuli were presented in a block design with 20 s of a picture naming task alternating with 14 s of scrambled images as the control. Analysis of fMRI data was done using AFNI. Preprocessing included registration to the preoperative anatomical MRI and compensation for slice acquisition-dependent time shifts per volume. The temporal signal to noise ratio (tSNR) was computed for each voxel across the time series as the absolute value of the mean signal divided by its standard deviation and averaged across subjects.

### Voxel-Based Lesion-Symptom Mapping

Surgical lesions were manually labeled on postoperative MRI scans in AFNI. Lesion masks were then morphed to a normative space (MNI152) by first aligning anatomical postoperative scans to preoperative scans within each subject and subsequently warping preoperative scans to a template in MNI space using nonlinear registration in Advanced Normalization Tools Software (http://stnava.github.io/ANTs/). Once in standard space, lesions masks were verified and edited by an experienced neurosurgeon.

For the VLSM analysis, only voxels that were included in the lesion masks of at least 5 subjects were analyzed. In order to reflect relative changes in BNT scores following surgery, preoperative scores were regressed out of postoperative scores using an ordinary least squares linear regression model, and the corrected postoperative BNT scores were used as the behavioral measure of interest. A linear regression was used to correct postoperative scores as there was a linear relationship between preoperative and postoperative scores. Additionally, linear regression models are more robust and enable corrected scores to maintain a gaussian distribution.^29^ VLSM was implemented using a multivariate support vector regression (SVR) model in order to correlate the pattern of lesioned voxels to observed deficits using a multivariate lesion symptom mapping toolbox (https://github.com/atdemarco/svrlsmgui.git).^22^ The SVR model was implemented using an ε-insensitive SVM algorithm with a radial basis function kernel from the Statistics and Machine Learning Toolbox in MATLAB. Hyperparameters (cost, σ, and ε) were optimized using a Baysian optimization approach to minimize resubstitution loss. Permutation-based cluster level correction of resulting beta maps was done with 5,000 permutations and a p-value of 0.05 for voxel-level and subsequent cluster-level correction in order to correct for multiple comparisons.

Resection volume was not included as a covariate in this analysis. Given that resections were constrained to the left temporal lobe, there was less spatial variability across resections. Additionally, anterior temporal resections across patients differed primarily in the extent of the posterior margin of resection, which directly influences resection volume. Thus, in this analysis, removing the effects of resection volume would be expected to distort the relationship between resected voxels and observed deficits.

### ECoG Acquisition and Analysis

Of the 95 patients, 42 (24 females, 18-60 years) underwent intracranial EEG to localize seizure foci using subdural grid electrodes or depth stereo-electroencephalographic electrodes prior to resection. Subjects performed a picture naming task, in which they were presented with black and white line drawings of common objects and asked to name the object shown.^26,30^ Continuous audio recordings of each patient were carried out, and articulation times for each trial were manually labeled. ECoG data was collected with a sampling rate of 2,000 Hz and a bandwidth of 0.1-700 Hz using NeuroPort NSP (Blackrock Microsystems) or with a sampling rate of 1,000 Hz and a bandwidth of 0.15-300 Hz using Neurofax (Nihon Kohden). Electrode localization was performed via registration of postoperative CT with preoperative anatomical MRI scans.^31^ Only trials in which subjects responded correctly were used in the analysis. Analyses were done with trials time-locked to picture onset. The gamma band (65-115 Hz) analytic signal was extracted from the raw signal using a frequency domain bandpass Hilbert filter.^32,33,34,35^ Surface-based mixed-effects multilevel analysis (SB-MEMA) was used to estimate broadband gamma activity (BGA) at a population-level.^33,34,36,37,38^ SB-MEMA maps were calculated as a contrast in reference to the control condition to isolate regions where activity was greater for picture naming compared to scrambled images. Significance levels were computed at an alpha-level of 0.05 after using a family-wise error rate correction for multiple comparisons. VLSM results were colocalized to a standardized cortical surface in order to compute overlap between the cluster found in VLSM and SB-MEMA results. Additionally, electrodes were indexed to the closest node on the same standardized surface, and electrodes located within the region of overlap between the VLSM cluster and the thresholded SB-MEMA map were used to calculate group estimates of BGA traces.

## Results

### Resections

Anterior temporal lobectomy with amygdalohippocampectomy (ATL+AH) resections (n=36) included the majority of the hippocampus, amygdala, temporal pole, and parahippocampal gyrus (PHG) as well as variable portions of the fusiform gyrus, the inferior temporal gyrus (ITG), the middle temporal gyrus (MTG), and the superior temporal gyrus (STG). Anterior temporal lobectomy (ATL) resections (n=9) were mainly confined to neocortical regions, which included the majority of the temporal pole and variable portions of the fusiform gyrus, ITG, MTG, and STG with relative sparing of the hippocampus, amygdala, and PHG. Additionally, 4 patients underwent modified anterior lobectomies. One patient underwent an ATL with amygdalotomy and relative sparing of the hippocampus. One patient underwent an ATL+AH with additional resection of portions of the insula. One patient underwent an ATL+AH with additional resection of a portion of the mesial frontal region. One patient underwent an ATL with additional resection of a portion of the basal frontal lobe. Resection overlap of all subject who underwent ATL procedures is shown in Supplementary Fig 1a. Laser interstitial thermal ablations (LITT) of the mesial temporal lobe (n=31) were confined to the hippocampus and amygdala and occasionally included a small portion of PHG immediately adjacent to the hippocampal region (Supplementary Fig 1b). Focal resections (n=15) were tailored based on individual clinical evaluations and were distributed across the left temporal lobe. Four patients had resections confined primarily to the temporal pole. Two patients underwent a laser ablation of a temporal periventricular nodular heterotopia (PVNH). Four patients underwent small resections located within the temporal-occipital cortex. One patient underwent a temporal mass resection involving lateral anterior temporal regions. One patient had a temporal topectomy in the lateral temporal area involving mostly MTG. One patient had a resection that included a portion of the anterior STG. One patient had a small resection located within posterior STG. One patient underwent an extensive ATL that included larger portions of the fusiform gyrus, ITG, and MTG. Resection overlap across all patients is shown in Fig 1. A threshold was applied to the resection overlap map to show only voxels resected in at least 5 patients, which reflects the voxels included in the VLSM analysis.

**Figure 1.**
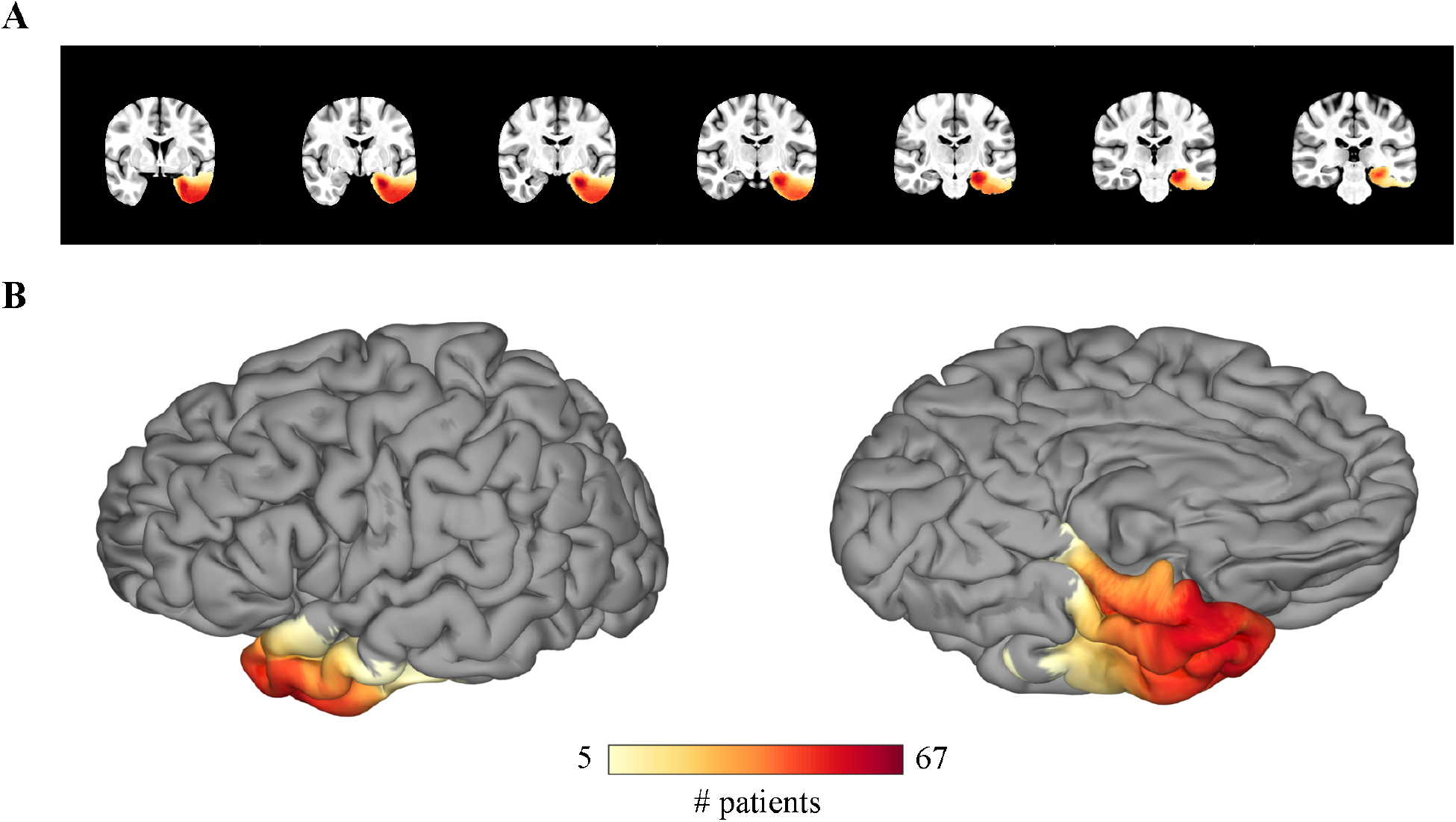
Lesion Overlap. The resection mask coverage across all subjects shown in the volume **(A)** and on the cortical surface **(B)** with a threshold applied to show only voxels that were included in the resections of at least 5 subjects.

### BNT Scores

The average raw preoperative BNT score was 45.35 +/- 8.85, and the average raw postoperative BNT score was 41.75 +/- 10.30 (Fig 2a). Prior to fitting the SVR model, the preoperative BNT score was regressed out of the postoperative BNT score using a linear regression. The resulting corrected postoperative BNT scores were represented as a standardized percentage centered at 0%, which corresponded to no change between preoperative and postoperative scores (Fig 2b). These standardized percentage scores can be interpretated as a single score representation of BNT performance following surgery relative to their preoperative performance. No change in BNT performance corresponded to 0% (grey dotted line in Fig 2b), a 5-point increase in BNT performance following surgery (the reliable change index for significant improvement) corresponded to 10.57% (green dotted line in Fig 2b), and a 4-point decrease in BNT performance (the reliable change index for significant decline) corresponded to -8.46% (red dotted line in Fig 2b).^39^ Fig 2c shows the correlation between preoperative and postoperative BNT scores both before and after linear regression. Scatter points are colored as clinically significant improvement (green), clinically significant decline (red), or clinically insignificant change (grey). The relationship between corrected postoperative BNT scores and the difference between postoperative and preoperative BNT scores is shown in Supplementary Fig 2.

**Figure 2.**
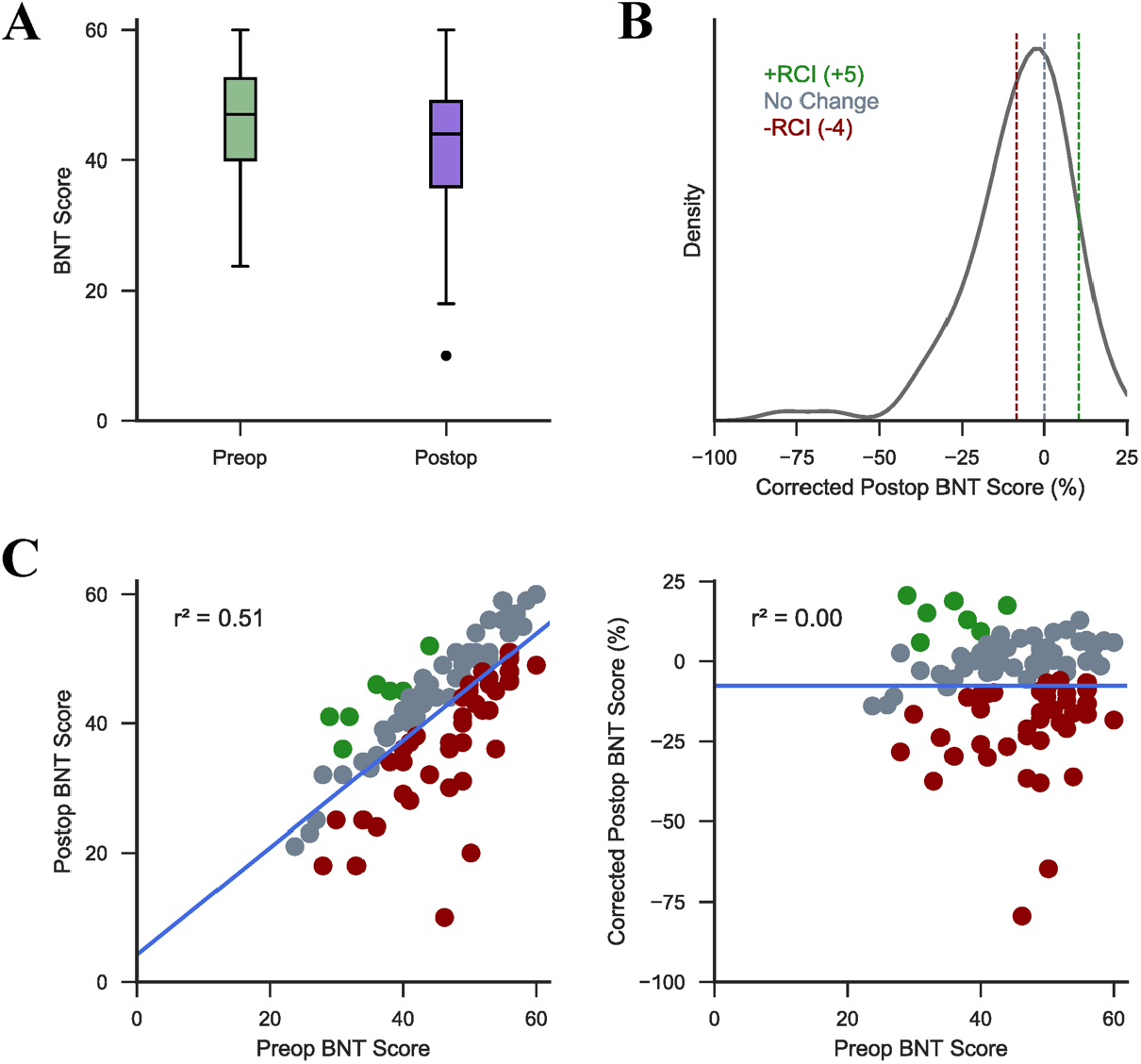
SVR Covariates. **(A)** The distribution of raw preoperative and postoperative BNT scores across all subjects. **(B)** The distribution of postoperative BNT scores following regression of preoperative BNT scores displayed as a standardized percentage. Dotted lines indicate corrected scores corresponding to a difference between postoperative and preoperative BNT scores of 0 (no change; grey), +5 (+RCI; green), and -4 (-RCI; red). **(C)** The correlation between preoperative and postoperative BNT scores (displayed as the percentage of correct answers) before (left) and after (right) regression. Scatter points are colored as clinically significant improvement (green), clinically significant decline (red), or clinically insignificant change (grey).

Statistical analyses were performed to examine the effect of language dominance on naming decline following left temporal lobe resections using unpaired, two-sample t-tests. In patients who underwent left ATL procedures, the change in BNT performance following surgery was significantly different in left language dominant patients compared to right language dominant patients (two-tailed p < 0.001) with left language dominant patients exhibiting a greater decline following resection (one-tailed p < 0.001). In patients who underwent left LITT procedures, there was no difference in the change in BNT performance following surgery between left and right language dominant patients (two-tailed p = 0.53).

### VLSM

Fig 3a and 3b show the unthresholded Beta map computed after fitting the multivariate SVR model. The Beta value of a given voxel represents its contribution to BNT performance where negative values indicate a decline in BNT performance following resection of the corresponding voxel. The Beta map revealed that resection of posterior inferior temporal regions contribute more to a decline in naming compared to anterolateral temporal regions. Fig 3c and 3d show the most significant cluster contributing to decline in BNT performance following permutation-based cluster level correction of the Beta map (p = 0.0248). The cluster primarily included the fusiform gyrus, PHG, and ITG. It had a volume of 5,656 mm^3^ and a center of mass located in the fusiform gyrus ([-33.1, -12.3, -33.4] in MNI space).

**Figure 3.**
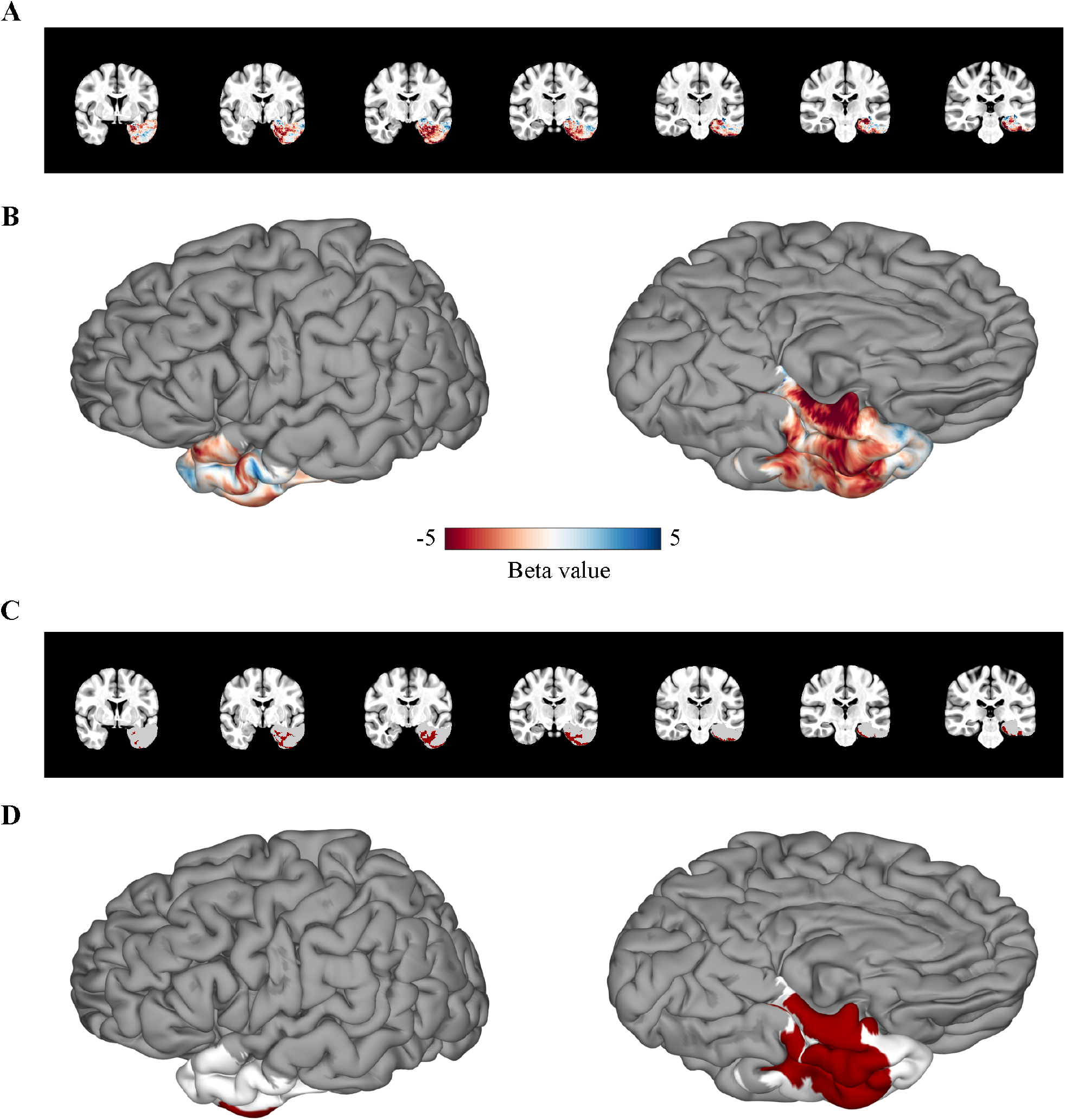
VLSM Results. **(A-B)** The Beta map computed using a multivariate SVR model shown in the volume **(A)** and on the cortical surface **(B)**. More negative values indicate that resection of the given voxel results in decline in BNT performance. **(C-D)** The most significant cluster following permutation-based cluster level correction of the Beta map shown in the volume **(C)** and on the cortical surface **(D)**. Regions included in the cluster are shown in red, and regions not included in the cluster but included in the analysis are shown in white.

Less than 3% of voxels within the significant cluster were located within the hippocampus. To look more closely at the role of the hippocampus and its effect in naming, we computed the correlation between the percent of the hippocampus removed and the corrected postoperative BNT score (r = -0.21, p = 0.044). We found that the percentage of the hippocampus removed was not strongly correlated with naming decline and explained less than 5% of the variance in corrected BNT scores (Supplementary Fig 3).

To investigate the nature of the relationship between the posterior extent of ATL resections and decline in BNT performance, we computed the correlation between the distance from the tip of the temporal pole to the most posterior coordinate of the resection mask and the corrected postoperative BNT score across subjects who underwent standard ATL or temporal pole resections (fig 4). We found that the extent of the posterior margin of the resection was linearly correlated to a decline in postoperative naming (r = -0.58, p < 0.001), with a significant decline in BNT performance (RCI = -4) occurring once the posterior margin reached ∼6 cm from the tip of the temporal lobe.

**Figure 4.**
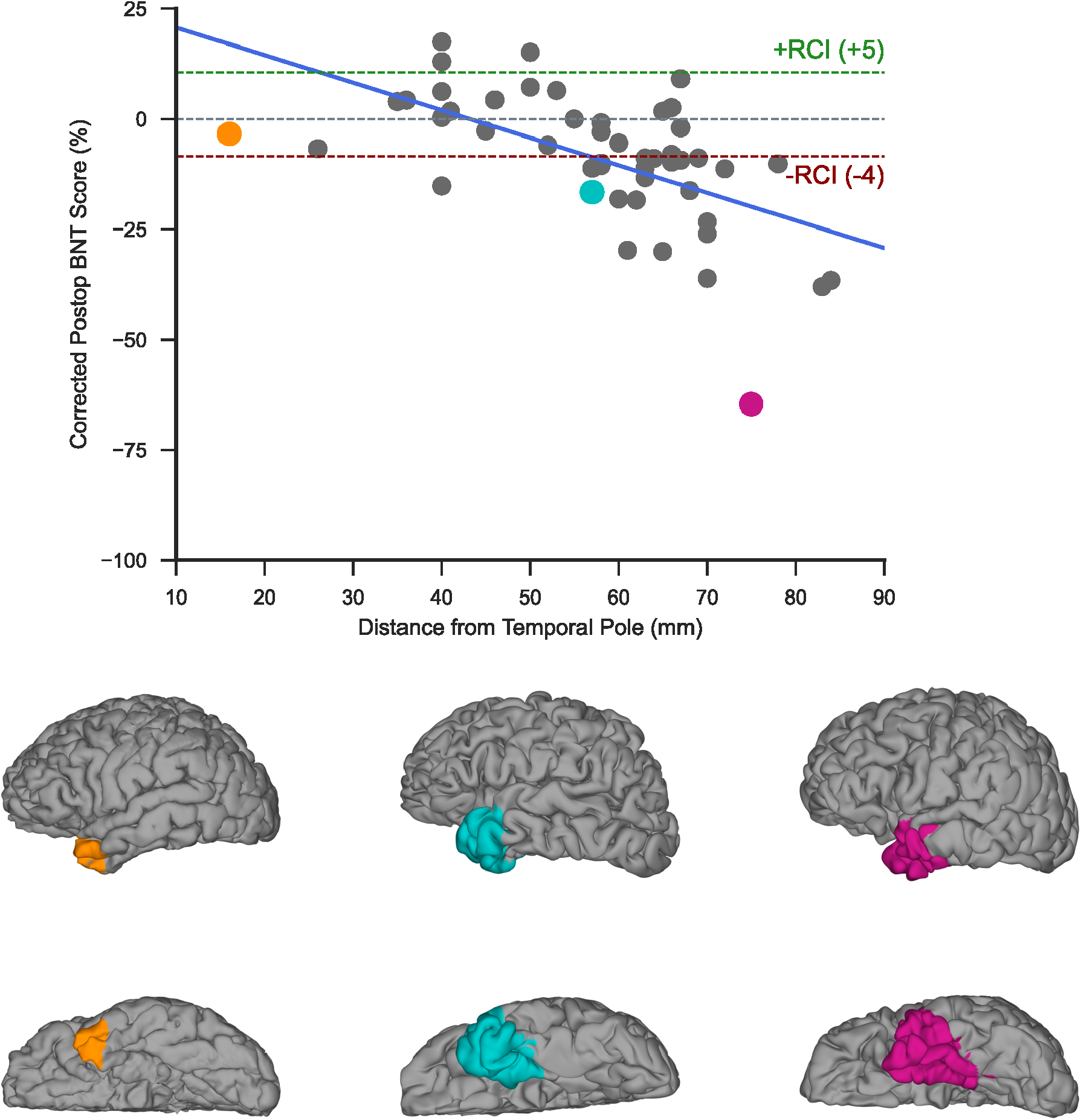
Analysis of Posterior Extent of ATL Resections. The correlation between corrected BNT score and the posterior margin of ATL resections measured as distance from the tip of the temporal pole (r = -0.58, p < 0.001). The resections of three examples patients are shown on the cortical surface and highlighted in the scatter plot in the corresponding color.

The average tSNR values per voxel of fMRI scans acquired during a picture naming task were assessed in a subset of patients from our cohort (fig 5). Regions with the lowest tSNR included the orbitofrontal cortex and the ventral temporal cortex. Whole brain tSNR ranged from 2.35 to 131.5 with an average of 79.40 +/- 21.73. The mean tSNR of the significant cluster computed using VLSM was 51.93 +/- 21.29.

**Figure 5.**
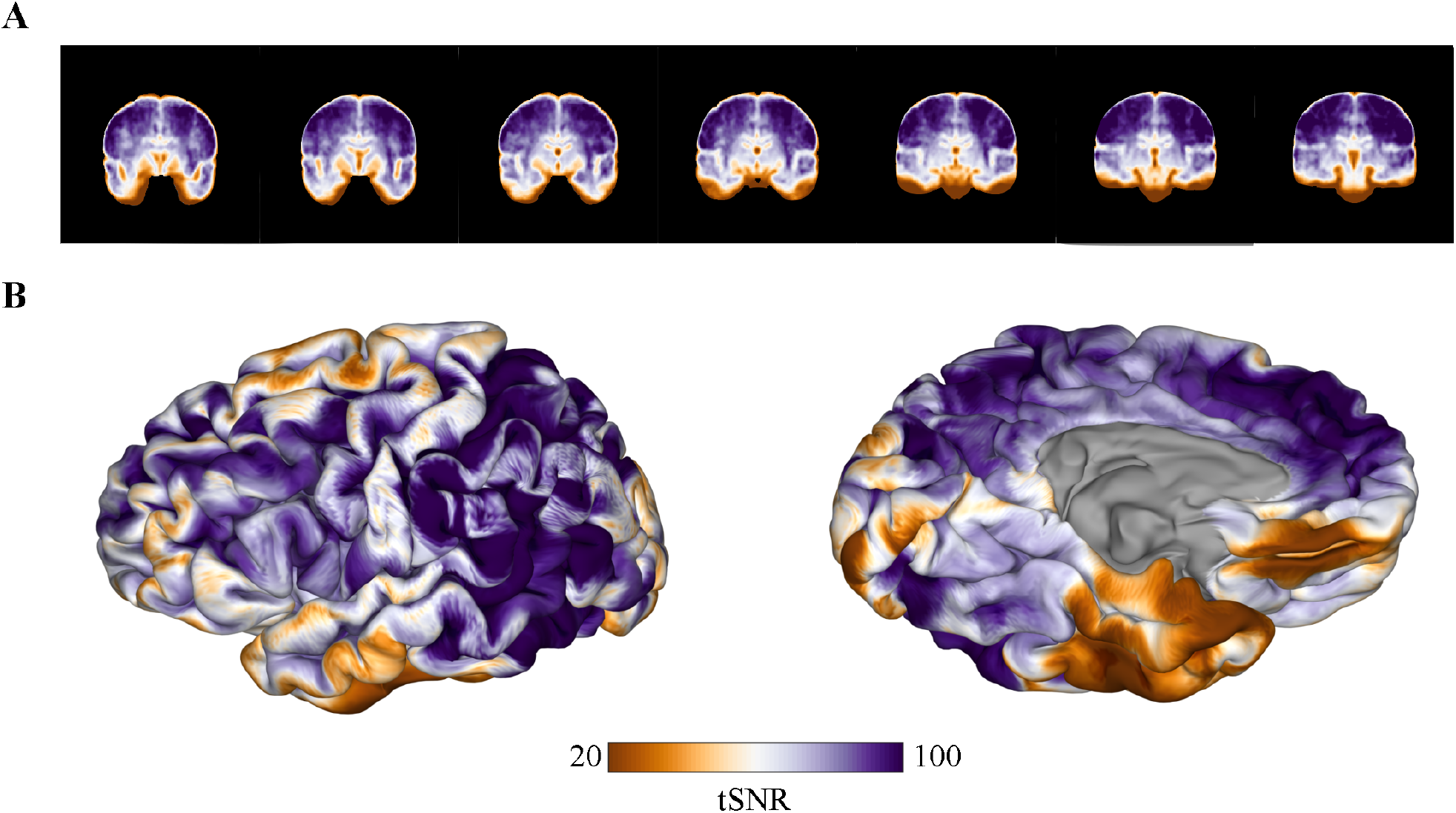
fMRI tSNR. The average tSNR calculated on fMRI scans acquired during a picture naming task shown in the volume **(A)** and on the cortical surface **(B)**.

### ECoG Analysis

The average reaction time for picture naming was 1,355 ms with a mean accuracy of 93.65 +/- 0.04%. Fig 6a shows the electrode coverage across patients with ECoG included in the analysis. SB-MEMA analysis during picture naming showed peak BGA spreading anteriorly along the ventral temporal stream 500 to 750 ms following picture onset, and regions with greater cortical responses to picture naming compared to scrambled images were located primarily within the posterior ventral temporal lobe (fig 6b). This is concordant with prior work revealing a semantic-specific increase in activity in this region using both fMRI and ECoG.^40^ The thresholded SB-MEMA map overlapped with 78.8% of the significant cluster computed using VLSM (fig 6c). Overlap was primarily seen in the fusiform gyrus. Fig 6d-e shows the average percent change in BGA relative to picture onset during pictures (purple) and scrambled images (black) for electrodes located within the overlap between the SB-MEMA map and the VLSM cluster with an absolute BGA change of at least 25%. There was a significant increase in BGA compared to the control condition (scrambled images) beginning around 250 ms following picture onset with a peak increase occurring within 500 to 750 ms. Significance was computed using an unpaired t-test with a false discovery rate (FDR) corrected p-value of 0.05.

**Figure 6.**
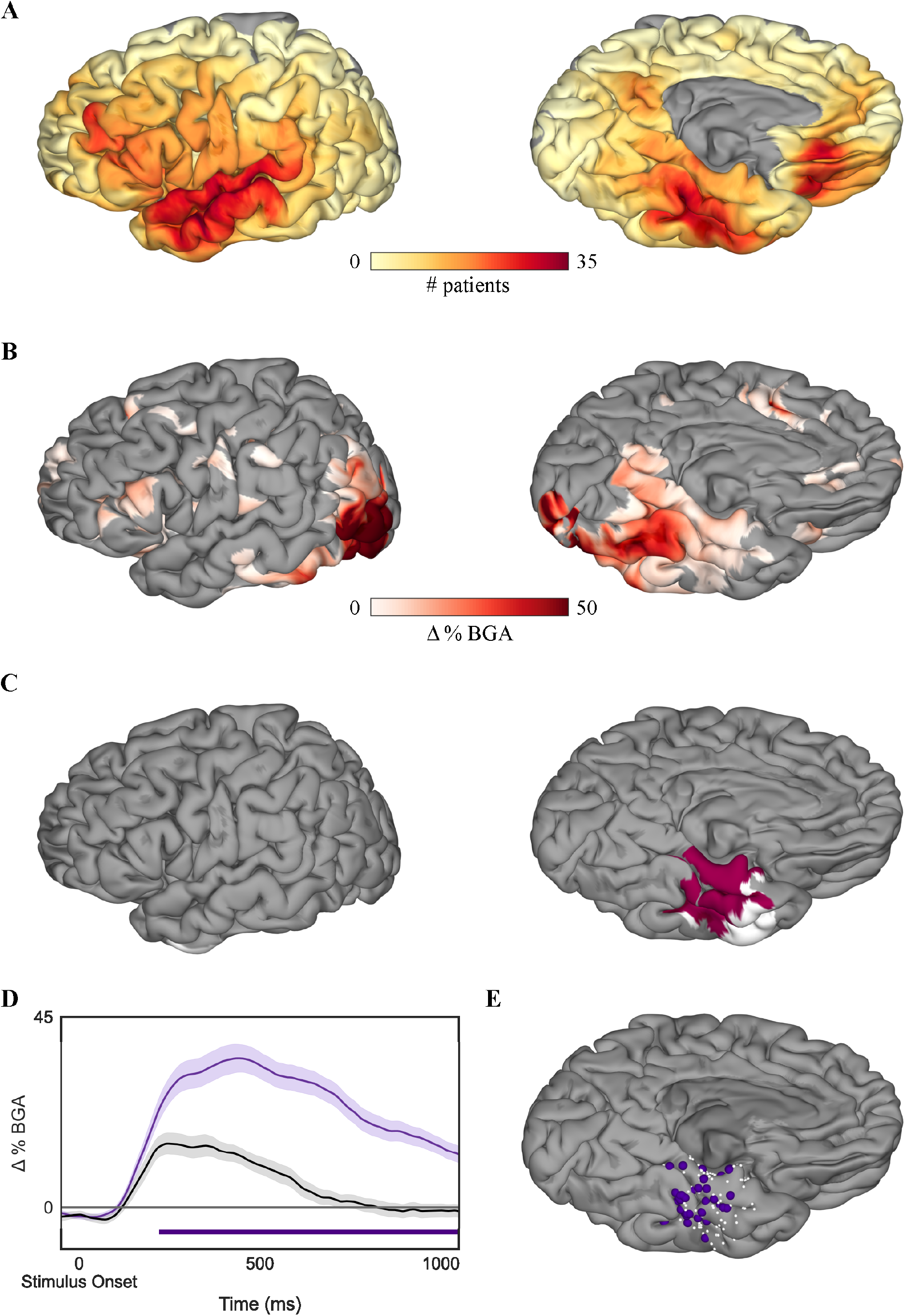
SB-MEMA Results. **(A)** Coverage of surface recording zones for all left hemisphere electrodes included in the study. **(B)** Surface-based group-level ECoG estimate of group BGA 500 to 750 ms following picture onset for pictures of objects > scrambled images. Maps are restricted to regions with significant activity (p < 0.05, corrected) and BGA change > 2.5%. **(C)** Regions in pink represent the overlap between active regions as determined by the thresholded SB-MEMA map and the VLSM significant cluster. Regions in white indicate nodes in the VLSM significant cluster that were not significantly active. **(D)** Time series average of group estimates of BGA percent change +/- 1 standard error of the mean following picture onset. Data are smoothed with a Savitsky-Golay filter (third order, 251 ms length). Significant increase from the control condition (scrambled images) is indicated by the horizontal bar (unpaired t-test, p < 0.05, FDR corrected). **(D)** Electrodes located within the VLSM cluster and SB-MEMA overlap. Electrodes with significant a cortical response included in the average trace are shown in purple, and electrodes that did not have a significant cortical response are shown in white.

## Discussion

We isolated the critical cortical constituents for confrontation naming using multivariate VLSM and integrated these findings with ECoG within the same patient population to derive quantifiable measures of convergence between lesion-deficit localization and brain activity. We found that damage to the dominant posterior ventral temporal cortex (pVTC) is significantly associated with pervasive deficits in visual naming. Together, these data firmly establish the role of the fusiform gyrus as a critical semantic access hub that is essential for lexical retrieval.^40,41,42^

Previous studies using cortical stimulation mapping have highlighted the importance of pVTC, also referred to as the basal temporal language area (BTLA), in confrontation naming.^40,43^ More specifically, it has been shown that disruption of the fusiform gyrus, PHG, and ITG results in speech arrest.^43,44^ Additionally, Forseth *et al*.^40^ observed that stimulation of the fusiform gyrus specifically disrupted object naming for both pictures and auditory descriptions without disruption of sentence repetition or sensorimotor effects, which implicates its role in semantic processing as opposed to audio-visual integration. Despite clear evidence for acute disruption in language function following electrical stimulation, previous studies have underplayed the role of this region in chronic lesions, reporting that resection of the BTLA results in no significant language impairments or only transient aphasias with the majority of language deficits resolving within one month following surgery.^43,45,46^ It has been hypothesized that these findings might suggest that the BTLA does not play an important role in language^47,48,49^ or that its role within language networks is not essential.^49,50^ However, other studies have found BTLA resections to be associated with more pervasive aphasias.^16,51,52^ Our data show that resections involving the left dominant BTLA do result in long-term language deficits with many patients reporting noticeable word finding difficulties at the time of postoperative testing, which took place at least 3 months following surgery. Furthermore, we found that left language dominant patients experienced significantly greater declines in naming ability compared to right language dominant patients following left ATL procedures, indicating that the function of the dominant BTLA is distinct from that of the nondominant BTLA.

Despite accumulating evidence implicating pVTC in semantic processing, functional neuroimaging studies have reported variable results regarding semantic-associated activations in the ventral temporal lobe, and many have localized semantic function to more lateral temporal regions.^53,54,55,56^ However, limitations of echo planar imaging may restrict the ability to study the medial temporal lobe using fMRI. Due to its proximity to both the sinuses and the skull, this area is particularly affected by image distortion and signal loss caused by susceptibility-induced magnetic field gradients, and efforts to minimize these artifacts in order to improve accuracy often result in a reduced SNR.^57,58^ We found that ventral temporal regions, especially those identified as significant by our VLSM analysis, were associated with a low SNR on fMRI, which may explain why this region has been under-appreciated as the locus responsible for postoperative deficits.

The exact role of the hippocampus in naming is not clearly understood. It has been shown that laser ablation of the hippocampus and amygdala reduces the risk of postoperative naming impairment in comparison to standard ATL procedures.^52,59,60,61^ Concordantly, we did not find a significant association between the hippocampus and naming performance, and less than 3% of voxels included in the significant cluster belonged to the hippocampus. Additionally, the percentage of the hippocampus removed or ablated was not significantly correlated with a change in BNT performance. Thus, our results do not support that the hippocampus is an essential component of object naming.

It is well-known that dominant ATL resections increase the risk of postoperative naming deficits.^41,62,63,64^ Based on these findings and studies in patients with semantic dementia, it has also been hypothesized that the temporal pole functions as the critical access hub in lexical semantic processing.^65,66^ Our results show that resection of the temporal pole within ∼3 cm of the anterior tip of the temporal lobe is not significantly associated with postoperative naming deficits. However, we found that the posterior margin of ATL resections was linearly correlated with postoperative naming scores, with resections extending more posteriorly resulting in larger naming declines and a clinically significant decline occurring if the resection extended more than 6 cm from the temporal pole. Trimmel *et al*.^67^ found the left pVTC to be functionally connected to the left anterior STG and temporal pole, and it has been suggested that naming decline following dominant ATL resections may be the result of partial disconnections between the resected regions and pVTC.^19,67^ While more anterior regions may support other aspects of semantic processing, our data does not support that these regions are essential to lexical retrieval. As such, language declines following resection of more anterior portions of the left temporal lobe may be the result of increased disruption of ventral temporal connections following ATL resections as opposed to removal of critical lexical semantic processing regions, and observed visual naming decline may reflect the proportion of disruption to the antero-posterior-basal temporal language network.

Using both multivariate VLSM and ECoG, we have demonstrated the essential contribution of pVTC, specifically the middle fusiform cortex, to lexical access. We found that posterior inferior temporal regions, centered around the fusiform gyrus, were significantly associated with a decline in visual object naming ability following resection, and the majority of these regions were most active on ECoG during picture naming just prior to articulation. These findings build on our prior work^40^ as well as a recent lesion-deficit mapping study^52^ elucidating the role of the fusiform gyrus in naming. Altogether, these results support the importance of pVTC in the lexical semantic network, and further implicate the fusiform gyrus as a critical lexical sematic access hub.

## Abbreviations

ATL: anterior temporal lobectomy
BGA: broadband gamma activity
BNT: Boston Naming Test
BTLA: basal temporal language area
ECoG: electrocorticography
ITG: inferior temporal gyrus
MTG: middle temporal gyrus
PHG: parahippocampal gyrus
pVTC: posterior ventral temporal cortex
SB-MEMA: surface-based mixed effects multilevel analysis
STG: superior temporal gyrus
SVR: support vector regression
VLSM: voxel-based lesion symptom mapping

## Acknowledgements

We thank all the patients who participated in this study, laboratory members at the Tandon lab (Oscar Woolnough and Jessica Johnson), neurologists at the Texas Comprehensive Epilepsy Program who participated in the care of these patients, and all the nurses and technicians in the Epilepsy Monitoring Unit at Memorial Hermann Hospital who helped make this research possible.

