## Supplementary Material for "The critical role of the ventral temporal lobe in word retrieval"

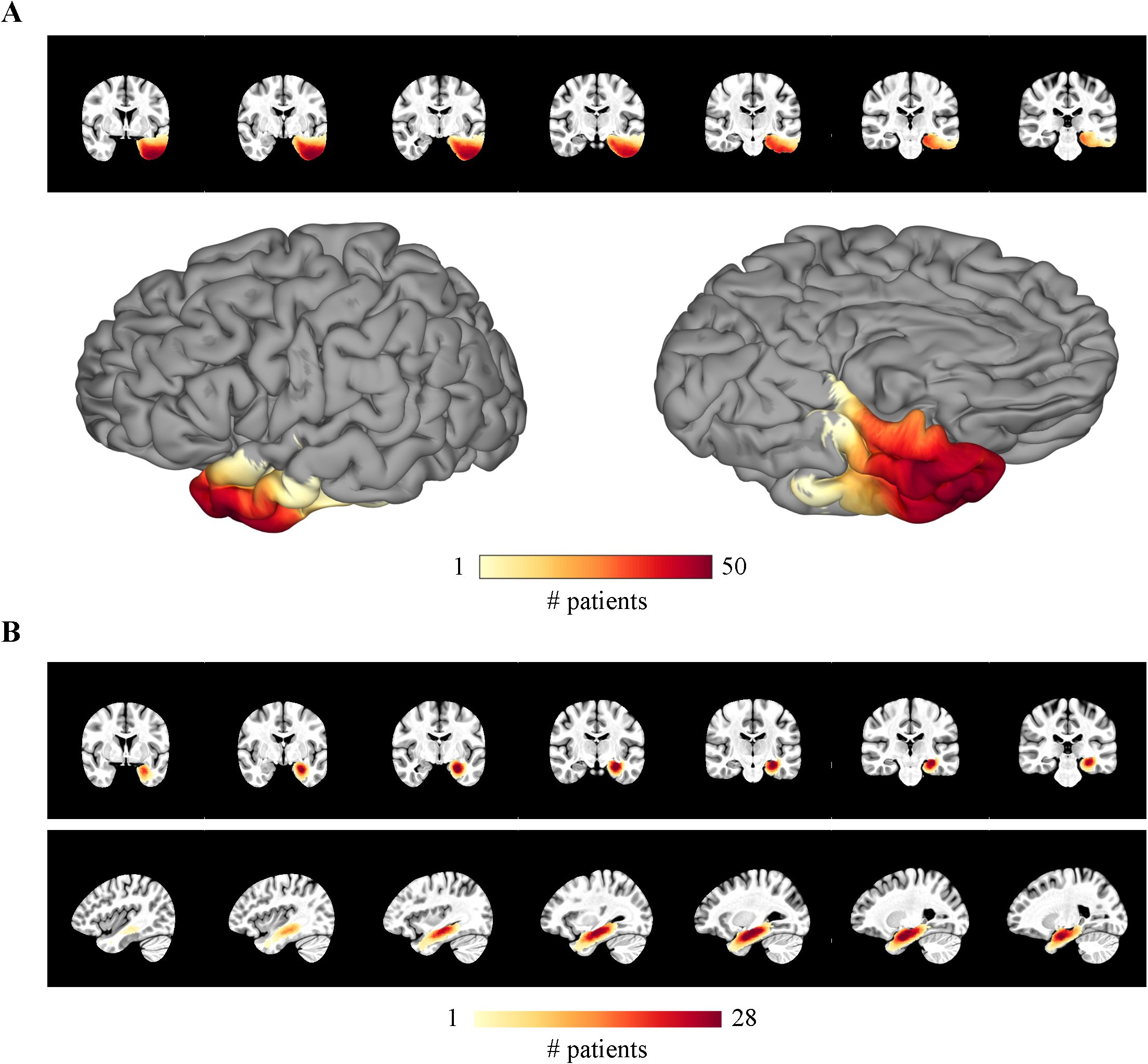

**Supplementary Figure 1. Lesion Overlap by Surgery.** The resection mask coverage across all subjects who underwent **(A)** a standard ATL procedure or **(B)** a selective laser ablation of the hippocampus and amygdala.

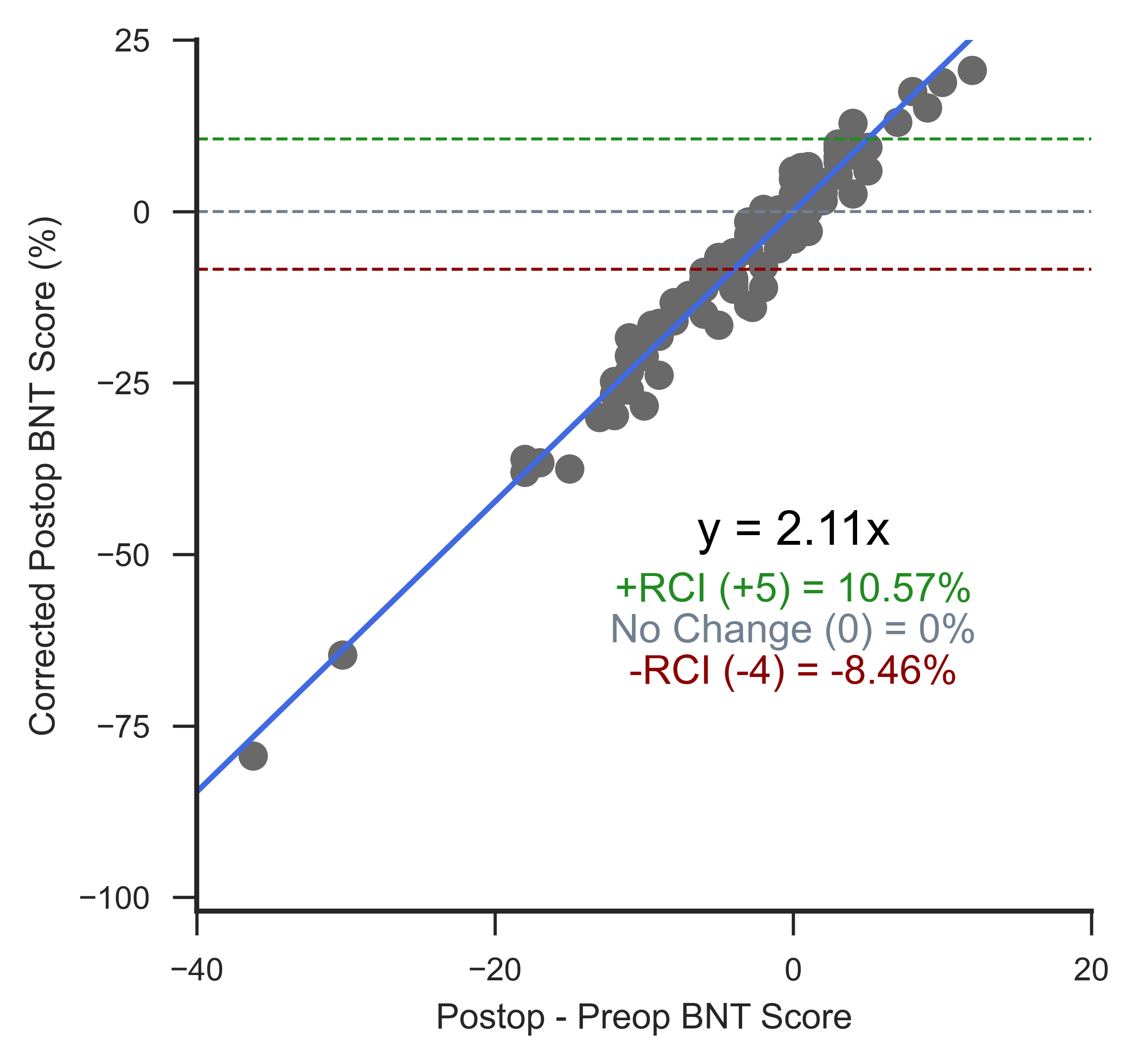

**Supplementary Figure 2. Corrected Postoperative BNT Score Interpretation.** The relationship between corrected postoperative BNT score and the difference between postoperative and preoperative BNT scores. Corrected postoperative BNT scores were centered at no change between preoperative and postoperative scores. An increase of 5 points postoperatively (significant improvement) corresponded to 10.57%; no change postoperatively corresponded to 0%; and a decrease of 4 points postoperatively (significant decline) corresponded to -8.46%.

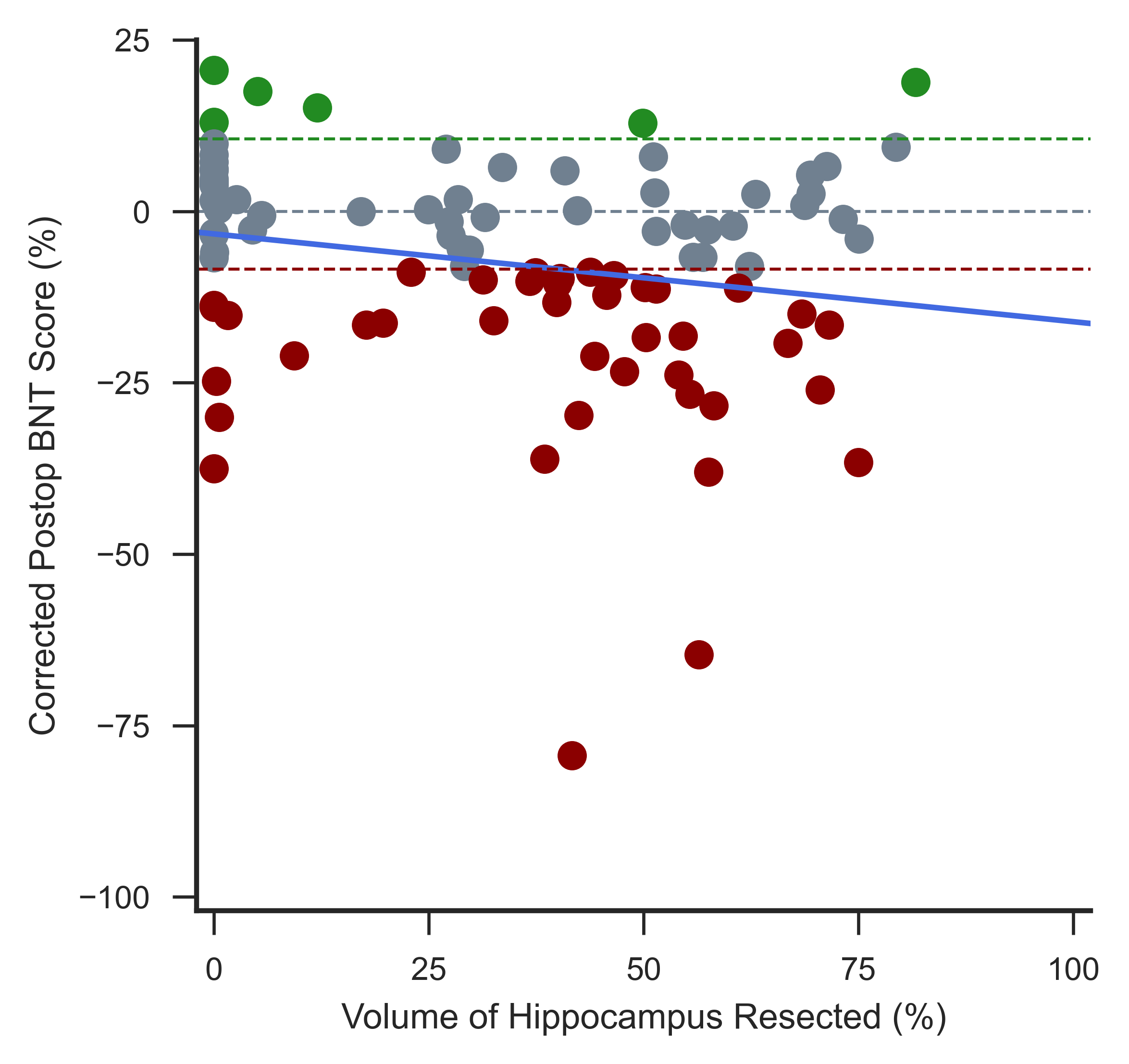

**Supplementary Figure 3. Hippocampal Resection and Naming Decline.** The correlation between the percentage of the hippocampus resected and corrected postoperative BNT score. There was not a strong correlation between the percentage of the hippocampus removed and naming decline (r^2^ = 0.04). Dotted lines indicate corrected scores corresponding to a difference between postoperative and preoperative BNT scores of 0 (no change; grey), +5 (+RCI; green), and -4 (-RCI; red), and scatter points are colored as clinically significant improvement (green), clinically significant decline (red), or clinically insignificant change (grey).
